# DNA hypomethylation during MSC chondrogenesis occurs predominantly at enhancer regions

**DOI:** 10.1101/625954

**Authors:** Matt J. Barter, Catherine Bui, Kathleen Cheung, Rodolfo Gómez, Andrew J. Skelton, Hannah R. Elliott, Louise N. Reynard, David A. Young

## Abstract

**Summary:** Regulation of transcription occurs in a cell type specific manner by epigenetic mechanisms including DNA methylation and histone modifications. Methylation changes during stem cell differentiation may play a key role in lineage specification. We sought to characterise DNA methylation changes during chondrogenesis of mesenchymal stem cells (MSCs) in order to further our understanding of epigenetic regulation in chondrocytes. The consequences of which has potential to improve cartilage generation for tissue engineering purposes and also to provide context for observed methylation changes in cartilage diseases such as osteoarthritis. We identified significant DNA hypomethylation during chondrogenesis including changes at many key cartilage gene loci. Importantly characterisation of significant CpG loci indicated their predominant localisation to enhancer regions. Comparison with adult cartilage and other tissue methylation profiles identified chondrocyte-specific regulatory regions. Taken together we have associated methylation at many CpGs with the chondrocyte phenotype.

**Abstract:** Regulation of transcription is determined in a cell type specific manner by epigenetic mechanisms including DNA methylation and histone modifications. Methylation changes during stem cell differentiation may play a role in lineage specification. Multipotent mesenchymal stem cell (MSC) differentiation into chondrocytes not only serves as a model for chondrocyte development but also provides an important source of cartilage for tissue engineering purposes. We sought to characterise DNA methylation changes during chondrogenesis to further understanding of epigenetic regulation but to also provide context for the changes identified during disease.

DNA cytosine methylation changes during human MSC differentiation into chondrocytes were measured by Infinium 450K methylation array. Methylation changes at gene loci were contrasted with gene expression changes. Chromatin states of significant methylation loci were interpreted by intersection with chondrogenesis histone modification ChIP-seq data. Chondrogenic and cartilage specific hypomethylation was utilised in order to identify a chondrocyte methylome. Articular cartilage and tissue panel DNA methylation was compared and alterations during osteoarthritis cartilage disease classified.

Significant DNA hypomethylation was identified following chondrogenic differentiation of MSCs including changes at many key cartilage gene loci. Highly upregulated genes during chondrogenesis were more likely to exhibit a reduction in DNA methylation. Characterisation of significant CpG loci indicated their predominant localisation in CpG poor regions which importantly are most likely to correspond to enhancer regions. Methylation level at certain CpGs following chondrogenesis corresponds to the level found in adult cartilage.

Taken together, considerable demethylation changes to the epigenetic landscape occur during MSC chondrogenesis especially at sites marked by enhancer modifications. Comparison with other tissues, including healthy and OA cartilage, associates CpGs to the chondrocyte phenotype and provides context for changes in disease.

## Introduction

Our skeleton acts as an essential framework for the overall structure of our body. Accurate generation of this frame is dependent upon carefully controlled chondrocyte differentiation and cartilage formation, processes that underpin the development of the long and short bones of the skeleton (Berendsen and Olsen, 2015). Chondrocytes, the sole cartilage cell type, develop from condensations of mesenchymal progenitor cells in a process known as chondrogenesis (Onyekwelu *et al.,* 2009). During chondrogenesis progenitors commit to a tightly coordinated chondrocyte differentiation program determined by temporal and spatial expression of multiple growth factors and transcription factors (Akiyama, 2008).

Epigenetic mechanisms, such as DNA methylation and histone modifications, provide cell-type specific regulation of gene expression essential for differentiation and maintenance of cell phenotype (Bernstein *et al.*, 2007). DNA methylation is a reversible process catalysed by DNA methyltransferases (DNMTs) on the fifth carbon of cytosine residues at CpG dinucleotides to form 5-methylcytosine (5mC) (Luo *et al.*, 2018). DNA methylation at gene promoter or enhancer sequences is frequently associated with gene repression where it correlates with the presence of inhibitory histone modifications and prevents the binding of transcription factors (Jones, 2012). Loss of DNA methylation occurs passively during cell replication or by conversion of 5mC back to cytosine via oxidised intermediates with the assistance of ten eleven translocation (TET) proteins (Wu and Zhang, 2014).

During development and cell differentiation DNA methylation is dynamic, correlating with changes in gene expression (Wiench *et al.*, 2011; Smith and Meissner, 2013). The role, whereabouts and dynamics of DNA methylation during chondrogenesis remains poorly understood. In chondrocytes the regulation of genes such as MMP13, IL1, iNOS, chondromodulin, collagen 9 and GDF5 is influenced by DNA methylation at specific CpGs (Aoyama *et al.*, 2004; Bui *et al.*, 2012; de Andres *et al.*, 2013; Hashimoto *et al.*, 2013; Imagawa *et al.*, 2014; Reynard *et al.*, 2014). Similarly DNA methylation may regulate gene expression during chondrogenesis where COLl0Al induction correlates with its promoter demethylation, and intereference with DNA methylation during chondrogenesis by treatment with 5-aza-C alters gene expression (Zimmermann *et al.*, 2008; El-Serafi *et al.*, 2011). While addition of methylation by DNMT3A has been found to regulate SOX9 expression in limb bud mesenchymal cells, and by DNMT3B to regulate cartilage metabolism and homeostasis (Kumar and Lassar, 2014; Shen *et al.*, 2017).

At the genome-wide level histone modifications are regulated during chondrogenesis and found to correlate with gene expression (Herlofsen *et al.*, 2013). In contrast DNA methylation at gene promoters, assessed by reduced representation bisulfite sequencing (RBBS), did not correlate with gene expression (Herlofsen *et al.*, 2013). Utilising the Infinium 450K methylation array the effect of ageing on cartilage DNA methylation and the similarity between MSC-derived cartilage methylation in comparison with cartilage engineered from articular chondrocytes have been studied (Bomer *et al.*, 2016; Peffers *et al.*, 2016).

Herein we focus directly on the differentially methylated CpGs during chondrogenesis in order to better define the chondrocyte methylome. Methylation changes are contrasted with gene expression to infer causality, while chromatin state information is integrated in order to further characterise regions of dynamic methylation. Further, a chondrocyte-specific methylation profile is established by comparison with cartilage and non-cartilage tissue methylation profiles, and the alterations in DNA methylation in osteoarthritis correlated. Interpretation of the epigenetic changes during chondrogenesis can provide context for those seen during deterioration of chondrocyte function in diseases such as osteoarthritis and also improve understanding of the processes governing differentiation of MSCs into chondrocytes for tissue regeneration purposes.

## Results

Human MSCs were differentiated into chondrocytes by culture in chondrogenic differentiation medium for 14 days in a scaffold-free Transwell insert. A cartilage disc is formed with a homogenous extracellular matrix and concomitant rapid upregulation of chondrocyte gene expression which is highly reproducible between different MSC donors (Murdoch *et al.*, 2007; Barter *et al.*, 2015). In order to identify CpG methylation changes during MSC chondrogenic differentiation a DNA methylation profile was generated with the Infinium HumanMethylation450 BeadChip. There was no change in median methylation level of all CpG sites between MSCs (Day0) and MSC-derived chondrocytes (Day14). However, differential methylation analysis identified 5950 differentially methylated CpG loci (DMLs) with a significant, greater than 10%, change in methylation level at Day14 compared with Day0 (Figure 1A and Supplementary Table 1). The vast majority of these (5802 DMLs) become hypomethylated during chondrogenesis, and the extent of methylation change is greater for CpGs becoming hypomethylated compared to hypermethylated (Figure 1B). A number of DMLs are found at key cartilage gene loci such as ACAN and SOX9 (Figure 1C).

**Figure 1.**
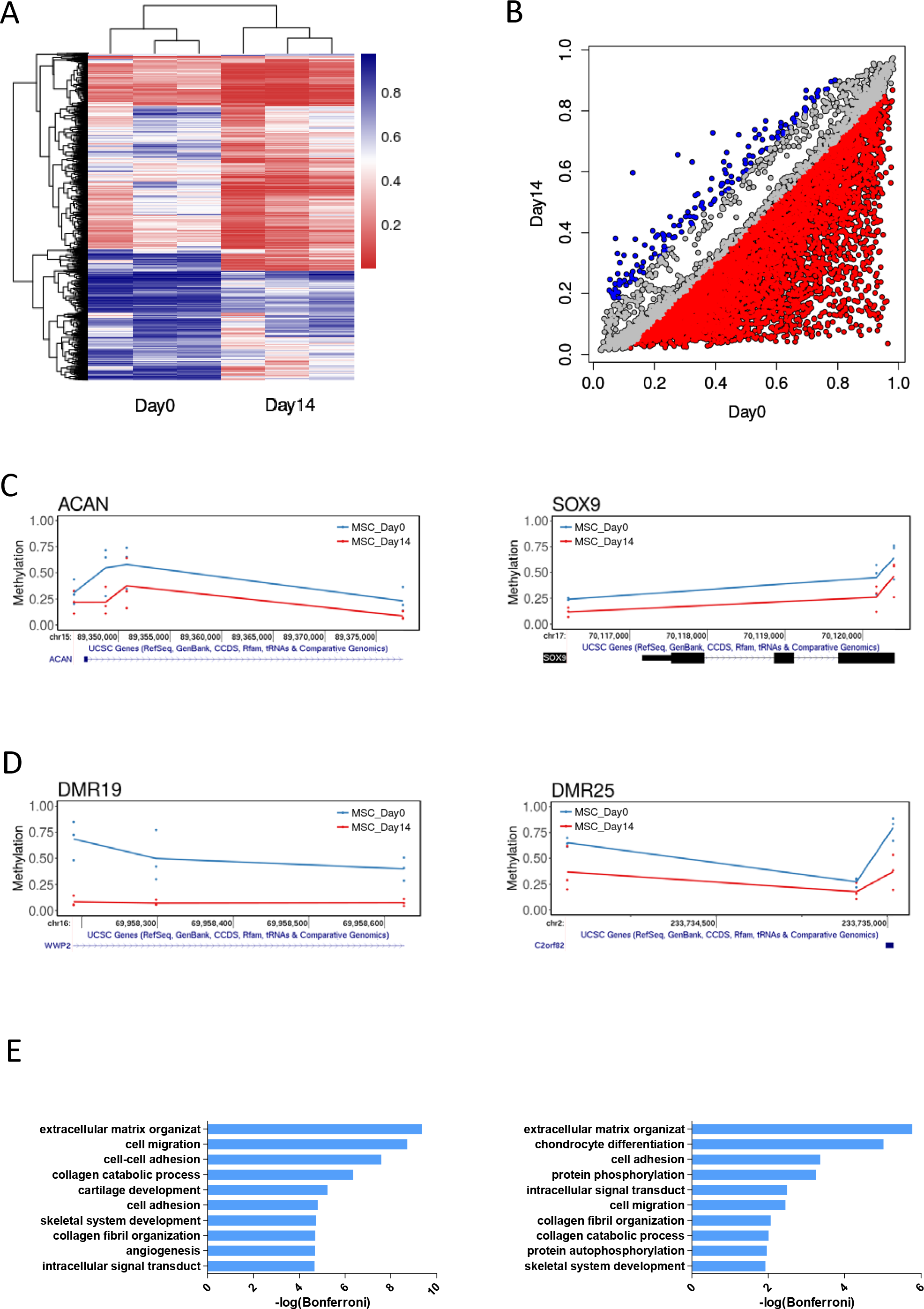
Differentially methylated CpGs during MSC chondrogenesis. A. Heatmap of significant CpGs in MSC Day0 and Dayl4. B. Correlation plot of significant CpGs Beta values at Day0 and Dayl4. Red indicates >10% hypomethylation, blue >10% hypermethylation at Dayl4. C-D: Genomic vignette of (C) significant CpGs at COL2A1 and SOX9 loci, (D) DMR6 at C2orf82 and DMR12 at WWP2 loci. Each donor is represented by a single dot and the average methylation for each timepoint represented by a line. Blue for MSC Day0, red for MSC Dayl4.

Grouping of DMLs in differentially methylated regions (DMRs) are considered an indication of regions with the potential to regulate gene transcription (Rakyan *et al.*, 2011). 1276 DMRs are found during MSC chondrogenesis (Supplementary Table 2), including a DMR found at the promoter of SNORC (C2orf82), the transcript of which we found previously as the most upregulated during MSC chondrogenesis, and a DMR in WWP2 a key cartilage development protein (Figure 1D) (Chantry, 2011; Barter *et al.*, 2015). GO term analysis of the genes with which these DMLs and DMRs are associated identifies terms consistent with the differentiation of cells into chondrocytes (Figure 1E).

We previously identified expression changes in greater than 2000 genes during MSC chondrogenesis (Barter *et al.*, 2015). Intersection of DNA methylation changes with gene expression changes identifies DMLs at 25% of upregulated genes and 15% of downregulated genes (Figure 2A). There is a greater enrichment of DMLs in more upregulated genes (Figure 2B).

**Figure 2.**
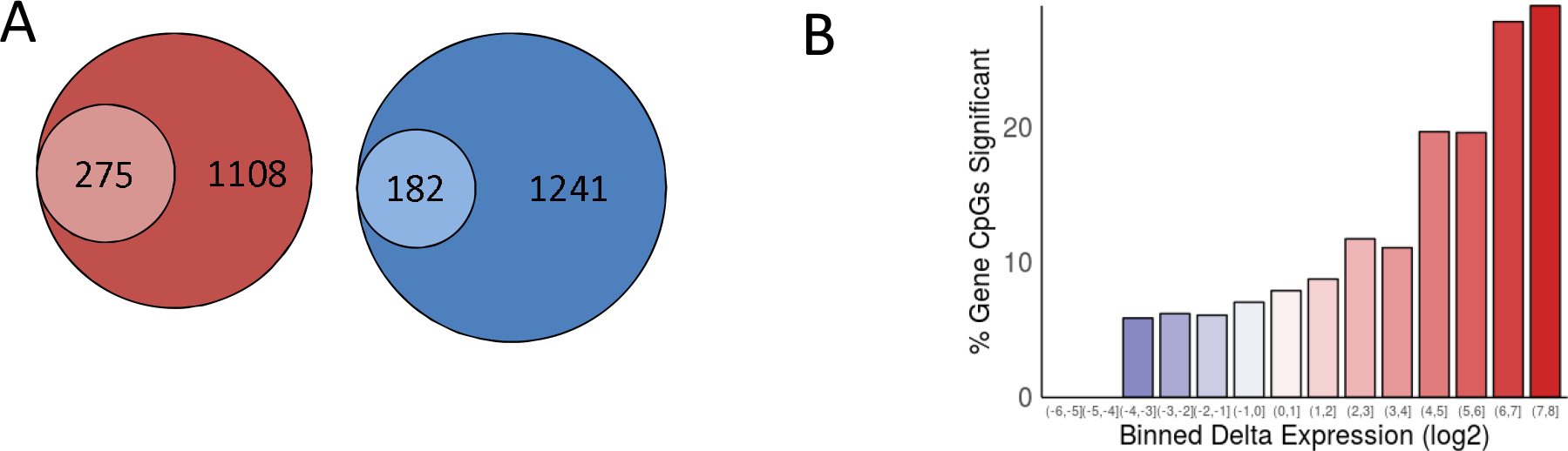
Comparison between methylation changes and gene expression changes. A. Proportion of differentially expressed genes during MSC chondrogenesis with methylation changes. Outer circle represent total 2fold up- (red) or down- (blue) regulated gene number in chondrogenesis. Inner circle represents number of genes with at least 1 significant CpG. B. Histogram showing enrichment of significant CpGs in comparison to direction and magnitude of binned log2 gene expression changes.

The context of the DMCs was further examined by examination of genomic features and chromatin states. In comparison with the distribution of probes across gene features in the array there is an enrichment of DMLs in gene body and intergenic regions (Figure 3A). These DMLs are also less likely to be found in or around CpG island areas. The hypomethylation occurs at all genomic features and locations (Figure 3B). A 15 chromatin state model of MSC-derived chondrocytes has previously been generated from the integration of multiple histone mark ChIP-seq data as part of the Roadmap Epigenomics project (Herlofsen *et al.*, 2013). CpG methylation levels during chondrogenesis were intersected with five functional categories of chromatin states from MSC-derived chondrocytes. Hypomethylation at DMLs was found at all chromatin states but was most extensive (>25%) for the enhancer state (Figure 3C). Taking all CpGs into account an empirical cumulative distribution frequency plot showed that the enhancer state exhibited the greatest hypomethylation during chondrogenesis (Figure 3D). Assessment of the distribution of DMLs across chromatin state categories indicates that a large proportion are found at enhancer states (Figure 3E).

**Figure 3.**
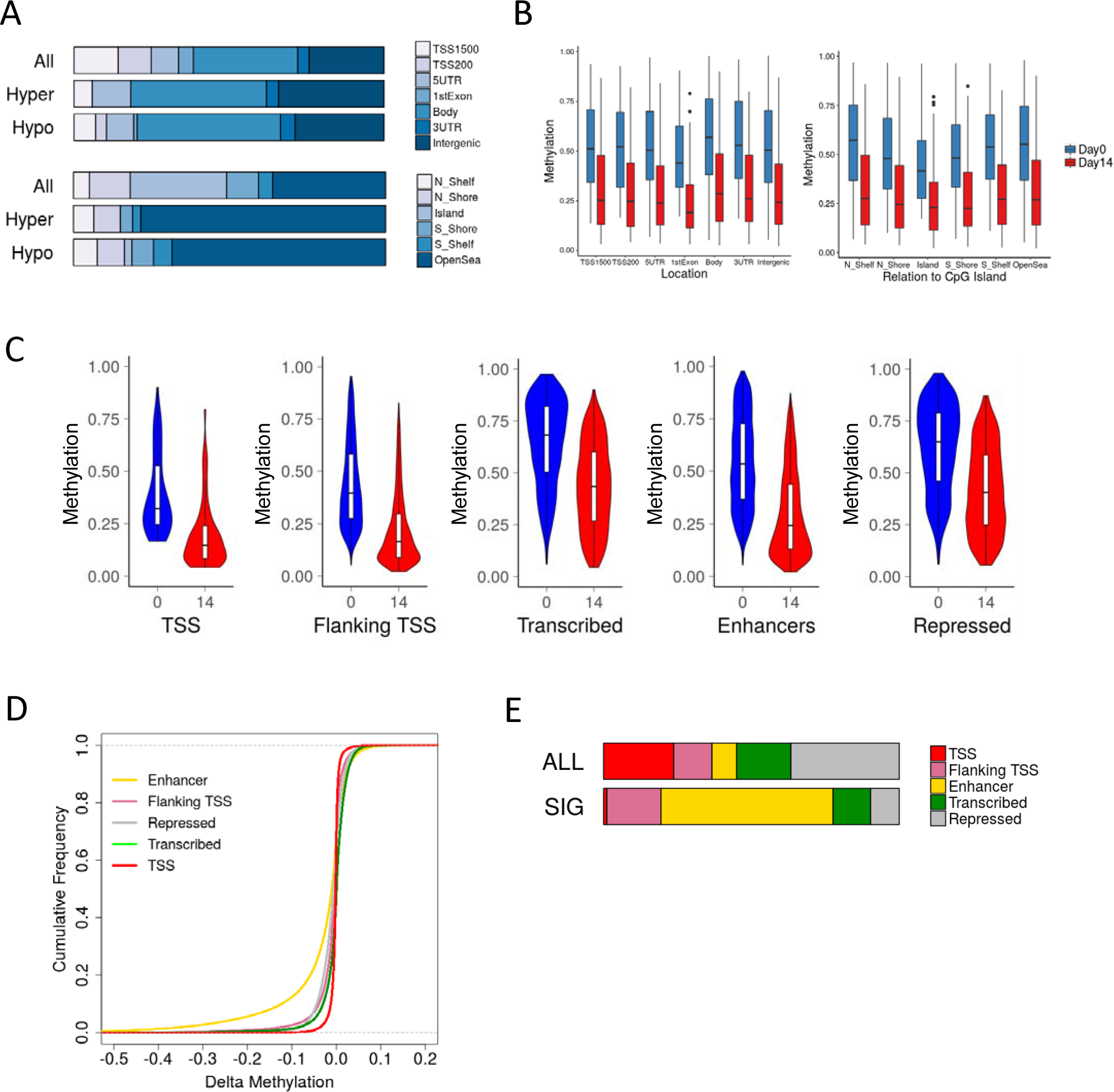
Genomic features and chromatin state of DMPs. A. Proportion of significant hypo- and hyper-methylated CpGs overlapping gene and CpG island features. The distribution of all CpG probes on the array are presented for comparison. B. Average methylation of significant hypo- and hyper-methylated CpGs overlapping gene and CpG island features in MSC Day0 and Dayl4. C-E. Roadmap Epigenomics project MSC-derived chondrocyte-specific chromatin states were collapsed into five functional categories. C. Average methylation of significant hypo- and hyper-methylated CpGs overlapping chromatin states in MSC Day0 and Dayl4. D. Cumulative frequency plot of methylation change at all CpGs during MSC chondrogenesis at each chromatin state. E. Proportion of significant hypo- and hyper-methylated CpGs overlapping chromatin states. The distribution of all CpG probes on the array are presented for comparison.

We previously identified DNA methylation levels in normal human hip chondrocytes from neck of femur fracture patients (GSE63695) (Rushton *et al.*, 2014). The final methylation level of DMLs during MSC chondrogenesis were compared with CpG methylation levels in healthy cartilage (NOF) and a number of other tissues with methylation data available from TCGA (https://cancergenome.nih.gov/). The level of methylation at DMLs at Day14 is more similar to the level of methylation in human articular chondrocytes (NOF) than Day0 (Figure 4A). In other tissues these DMLs have a higher level of methylation indicating a chondrocyte specific demethylation during chondrogenesis. For example DMLs at the CD109 and VGLL4 gene loci are methylated at levels similar to the other non-cartilage tissues at Day 0 but following chondrogenesis the methylation level is reduced to those found in articular chondrocytes (Figure 4B). To further characterise the chondrocyte DNA methylome tissue-specific methylation markers were sought. 1464 CpGs were identified with an average methylation beta value <0.5 in chondrocytes and >0.8 in >90% of tissues (Figure 4C). These CpGs are more highly methylated in MSCs before and after chondrogenesis. The COL11A2 and microRNA miR-140 gene loci include a number of CpGs exhibiting hypomethylation only in chondrocytes (Figure 4D).

**Figure 4.**
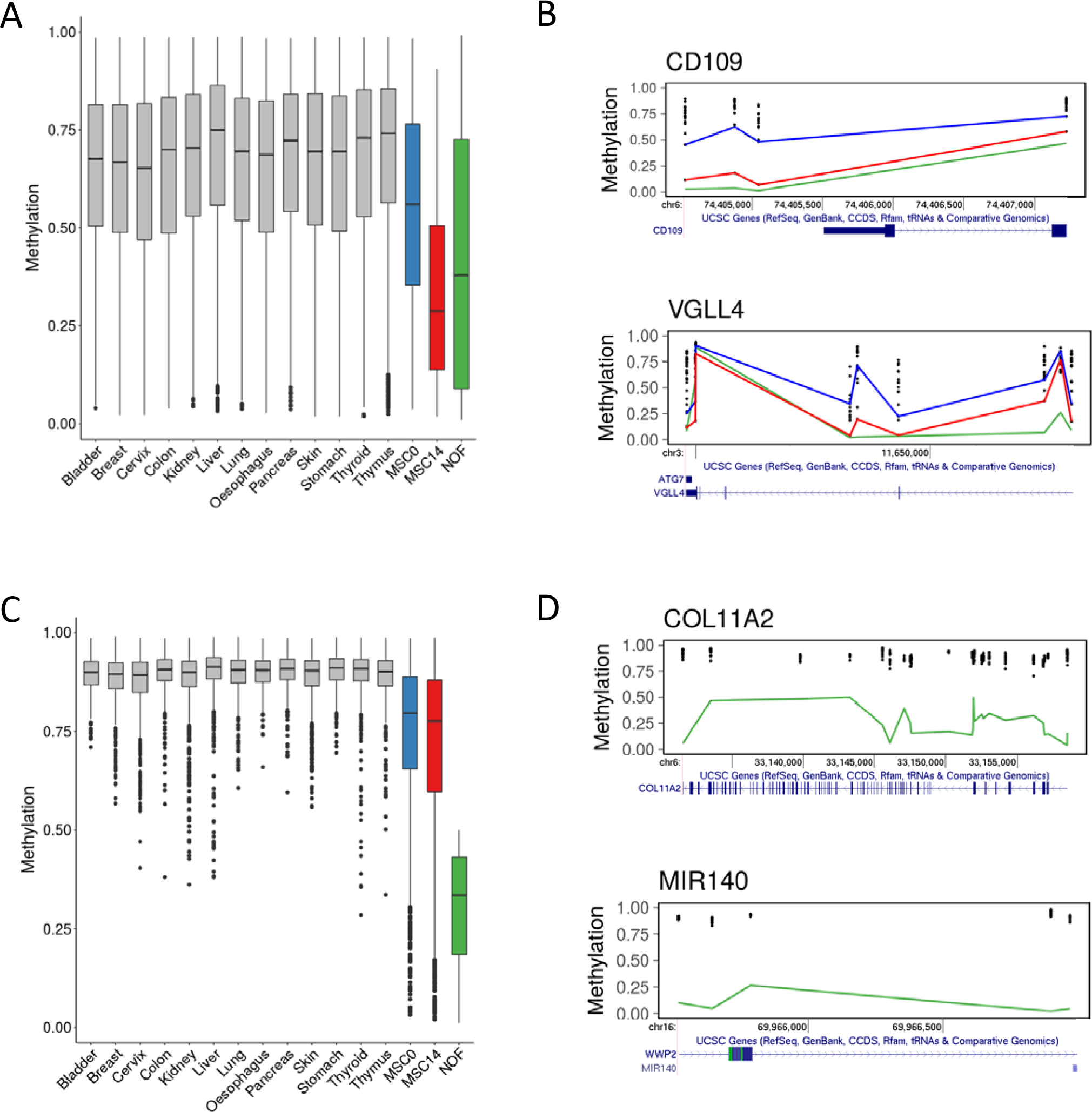
Methylation in chondrogenesis compared with human articular chondrocytes and other tissues. Mean methylation levels at Day0 and Dayl4, in human articular chondrocytes from neck of femur fracture patients (NOF), and in normal samples from a number of TCGA tissues were quantile normalised. A. Average methylation of significant CpGs during MSC chondrogenesis in comparison with NOF and other tissues. B. Genomic vignette of significant CpGs at VGLL4 and CD109 loci. C. Methylation level of 'cartilage-specific' hypomethylated CpGs in NOF compared with other tissues. D. Genomic vignette of 'cartilage-specific' hypomethylated CpGs at DIP2C and MIR140 loci. B & D. Each tissue is represented by a single dot. Lines indicate the methylation in MSC Day0 (blue), MSC Dayl4 (red), and NOF (green).

Changes in cartilage during OA may reflect a reinitiation of developmental pathways in the chondrocyte or a loss of chondrocyte phenotype due to disrupted epigenetic regulation (Loeser *et al.*, 2012; Reynard, 2017). Previously we identified DMLs between the hip cartilage of OA patients and healthy controls (Rushton *et al.*, 2014). A comparison between CpGs altered in OA hip cartilage and MSC chondrogenesis indicated a small, but greater than expected, proportion (~10%, hypergeometric distribution p<0.001) common to both (Figure 5A). For these 481 common CpG there is a reduction in methylation during OA consistent with the changes during chondrogenesis (Figure 5B). This is in contrast to the increase in mean methylation of all OA DMLs (Figure 5C). In comparison with the methylation of other tissues many CpGs altered in OA are found at a lower level of methylation in chondrocytes (Figure 5C).

**Figure 5.**
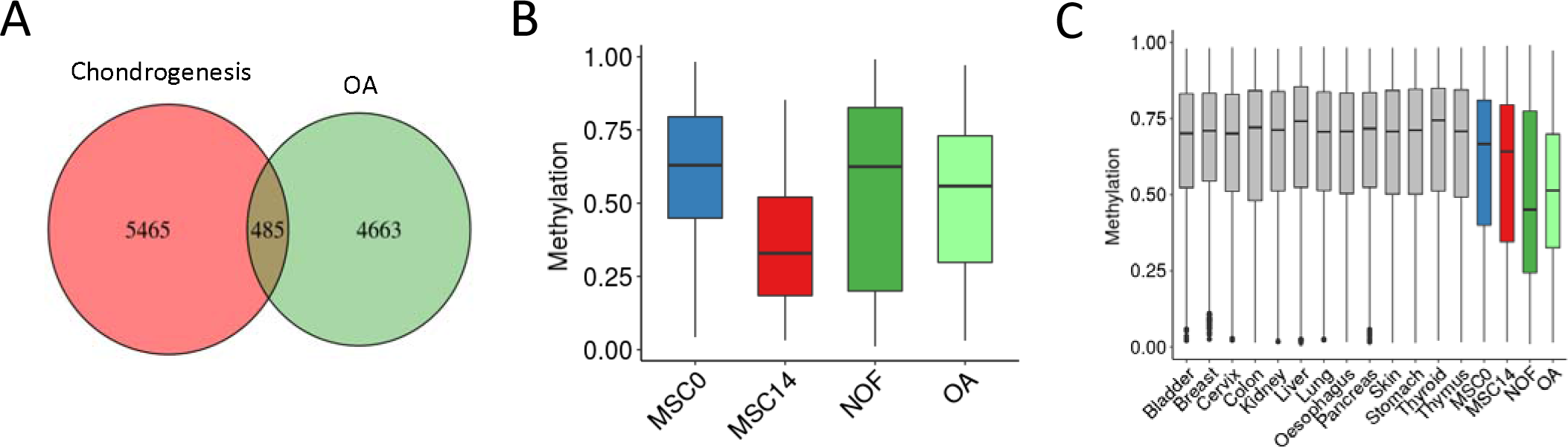
Overlap between DMPs in chondrogenesis and OA. A. Overlap between differentially methylated CpGs in MSC chondrogenesis and in hip OA. B. Average methylation of CpGs common to chondrogenesis and OA. C. Average methylation of all significantly differentially methylated CpGs in OA compared with other tissues.

## Discussion

DNA methylation establishes and reinforces cell-specific gene expression patterns during development and cell differentiation (Smith and Meissner, 2013). Here we show that DNA methylation changes, predominantly in the form of hypomethylation, are also a characteristic of differentiation of MSCs into chondrocytes during cartilage formation. These DMLs are enriched at enhancer regions consistent with the key role of cell-type specific enhancers in regulating gene expression. During MSC chondrogenesis the methylation level at significant CpGs becomes closer to the level found in articular chondrocytes, however significant differences remain between MSC-derived chondrocytes and adult chondrocytes.

By RBBS only limited changes in promoter methylation were identified during MSC chondrogenesis in a 3D alginate model (Herlofsen *et al.*, 2013), but more recently by Infinium 450K methylation array 1116 differentially methylated CpGs were identified across a timecourse of chondrogenic differentiation in a pellet model, albeit starting at Day14 (to Day49) in contrast to our comparison of undifferentiated vs. differentiated cells (Day0 vs. Day14) (Bomer *et al.*, 2016). The vast majority of differentially methylated CpGs during chondrogenesis herein become hypomethylated indicative of a loss of repression at these locations. Transcription factor binding can cause local demethylation (Edwards *et al.*, 2017) and thus the activation of the chondrocyte gene expression program might actively cause the loss of methylation. Consistent with this, hydroxymethylated cytosine (5hmC), a pathway intermediate generated during active demethylation, is found enriched at regulatory regions of genes during ATDC5 cell chondrogenic differentiation, although only a subset corresponded with changes in gene expression (Taylor *et al.*, 2016).

Interpretation of the genomic landscape and chromatin states harbouring significant CpGs indicated their enrichment in non-CpG rich enhancer regions of the genome. Consistent with the open-sea localisation of significant CpGs enhancers are mostly CpG-poor with variable methylation while in contrast 75% of promoters are within CpG islands with low methylation (Edwards *et al.*, 2017). Hundreds of thousands of enhancers exist in the human genome, potentially contributing the major function of non-coding DNA regions (Consortium, 2012). These cis-regulatory regions can drive transcription over long distance from their target gene, by for example forming chromatin loops, and offer both cell- and developmental-stage type specific regulation of gene expression (Long *et al.*, 2016). Cooperative action of transcription factors at enhancers and promoters facilitates chromatin access and activates gene expression (Long *et al.*, 2016). DNA methylation and DMRs at enhancers have been shown to correspond to enhancer activity of differentiation and cell-type specific genes (Wiench *et al.*, 2011). Numerous enhancers play key roles in regulating chondrocyte specific gene expression and skeletal development (Liu *et al.*, 2017). In particular both the regulation of SOX9 and the regulation by SOX9 are mediated by well-established enhancers (Ohba *et al.*, 2015; Yao *et al.*, 2015). During chondrogenesis the regulation of enhancer-associated H3K4me1 or H3K27ac modifications at distal cis-regulatory elements, up to 50kb from transcription start sites, correlates with gene expression suggesting that enhancer regions regulate many upregulated genes (Herlofsen *et al.*, 2013).

DMLs are enriched in genes associated with cartilage development GO terms, and grouping genes by the extent of their expression change during chondrogenesis indicated that the most upregulated genes contain more DMLs. However, many of the genes undergoing expression changes have no associated DNA methylation changes, and downregulated genes were also subject to hypomethylation. A number of factors may contribute to this including the baseline methylation level and chromatin state, the particular transcription factors regulating expression, as well as the genomic location of CpGs on array. Interestingly the converse is also true whereby binding of transcription factors to promoters even in the absence of active transcription can cause the local loss of DNA methylation (Edwards *et al.*, 2017). To better establish correlation whole genome methylation in combination with chondrocyte ATAC-seq data with gene expression and possible overlap of methylation and transcription factor binding sites (Liu *et al.*, 2018).

The disparity between MSC-derived cartilage and endogenous cartilage are well established owing to differences in lineage and developmental environment of the cells (Vinatier and Guicheux, 2016). At the epigenetic level the DNA methylation profiles are also quite distinct (Bomer *et al.*, 2016), however at many DMLs in chondrogenesis we identify a reduction in methylation level to the level found in adult articular chondrocytes, validating the utility of the MSC chondrogenesis model for chondrocyte differentiation. Notably these CpGs are found highly methylated in other tissues supporting the chondrogenesis-specific nature of the methylation changes. In particular the region containing WWP2 and miR-140 undergoes extensive demethylation during chondrogenesis and is found at highly methylated in all other tissues further cementing the cartilage-specific function of these genes (Chantry, 2011; Inui *et al.*, 2018). Genes with unexplored but possible functions in cartilage were also found hypomethylated in cartilage including CD109, a TGF-β co-receptor, and VGLL4 a regulator of Wnt signalling (Bizet *et al.*, 2011; Jiao *et al.*, 2017). COL10A1 expression … Previously methylation differences were also identified between MSC-derived cartilage from young and old donors however it is unclear whether these differences are due to baseline differences in the MSCs or as a result of differences in the differentiation capacity of the donors (Peffers *et al.*, 2016).

Providing context for the changes in OA we found a small overlap between methylation changes in MSC chondrogenesis and those CpGs altered in OA, with these DMLs again enriched in enhancers (Rushton *et al.*, 2014). Our data indicates that where there was previously no evidence of methylation changes in gene promoters during OA, such as ACAN and p21WAF1/CIP1, that dynamic methylation at enhancers regulating such genes should be considered (Poschl *et al.*, 2005; Sesselmann *et al.*, 2009). A number of studies have also identified DNA methylation changes in OA or damaged cartilage but it remains to be determined whether these correspond to enhancer regions (den Hollander *et al.*, 2014; Fernandez-Tajes *et al.*, 2014; Jeffries *et al.*, 2014; Alvarez-Garcia *et al.*, 2016; Bonin *et al.*, 2016; Zhang *et al.*, 2016; Steinberg *et al.*, 2017). The subset of chondrogenesis DMLs also significantly altered in OA become hypomethylated consistent with the cells reactivating developmental pathways (Loeser *et al.*, 2012; Reynard, 2017). However the majority of CpGs in OA become methylated and more similar to the level in other cell types indicative of a loss of chondrocyte identity. Synthesis of all OA DNA methylation studies by meta-analysis could provide a comprehensive catalogue of DMLs altered in disease for future comparison.

In conclusion, considerable demethylation changes to the epigenetic landscape occur during MSC chondrogenesis especially at sites marked by enhancer modifications. Comparison with other tissues, including healthy and OA cartilage, associates CpGs to the chondrocyte phenotype and provides context for changes in disease.

## Experimental Procedures

### Human Bone Marrow Stem Cell Culture

Human bone marrow mesenchymal stem cells (MSC) (from four donors, 18-25 years of age) were isolated from human bone marrow mononuclear cells (Lonza Biosciences, Berkshire, UK) by adherence over 24 hours to tissue culture plastic and were expanded in monolayer culture in Mesenchymal Stem Cell Growth Medium (Lonza) supplemented with 5ng/ml fibroblast growth factor-2 (R&D Systems, Abingdon, UK). Cultures were maintained in a humid atmosphere of 5% CO2/95% air at 37°C. Once cells reached confluence, they were passaged (P1) using Trypsin/EDTA at a split ratio of 1:3. Experiments were performed using cells between P2-P7. The phenotypes of all MSC donors were tested previously (Barter et al., 2015).

#### Chondrogenic Differentiation

MSC were resuspended in chondrogenic culture medium consisting of high glucose DMEM containing 100 μg/ml sodium pyruvate (Lonza), 10 ng/ml TGF-β3 (PeproTech, London, UK), 100 nM dexamethasone, 1x ITS-1 premix, 40 μg/ml proline, and 25 μg/ml ascorbate-2-phosphate (all from Sigma-Aldrich, Poole, UK). 5x105 MSC in 100μl medium were pipetted onto 6.5mm diameter, 0.4-μm pore size polycarbonate Transwell filters (Merck Millipore, Watford, UK), centrifuged in a 24-well plate (200g, 5 minutes), then 0.5 ml of chondrogenic culture medium added to the lower well as described (Murdoch et al., 2007). Media were replaced every 2 or 3 days up to 14 days. Donor 1 chondrogenesis was performed in triplicate, Donor 2, Donor 3 and Donor 4 in singlicate.

#### DNA isolation, bisulphite treatment and methylation array

Genomic DNA was extracted from cells prior to the induction of chondrogenesis (Day0) and from Day14 cartilagenous discs by disruption in Genomic Digestion Buffer using a small disposable plastic pestle and an aliquot of Molecular Grinding Resin (G-Biosciences, St. Louis, MO) followed by proteinase K digestion for 1 hour at 37°C then nuclei acid purification with Invitrogen PureLink Genomic DNA Kit according to manufacturer's instructions (Life Technologies, Paisley, UK). 1μg of genomic DNA was bisulphite converted using the EpiTect Bisulfite Kit (Qiagen). DNA methylation profiling of the samples was carried out by Cambridge Genomic Services (Cambridge, UK), using the illumina Infinium HumanMethylation450 Beadchip array (illumina, Inc., San Diego, USA).

#### Analysis of Infinium 450K methylation data

Methylation analysis was performed using the R/Bioconductor package Minfi. Raw IDAT files were read and preprocessed and probes were filtered for high detection p-value (P⍰>⍰0.05), location on sex chromosomes, ability to self-hybridize, and potential SNP contamination. Array normalization was carried out using the preprocessFunnorm function to generate M values for statistical testing and Beta values indicative of the percentage CpG methylation per probe. Differentially methylated probes were identified by fitting a paired linear model followed by statistical analysis using an empirical Bayes method (R/Bioconductor package Limma) then filtered by significance threshold (P⍰>⍰0.05, F-test, after correction for multiple testing using the Benjamini-Hochberg method). Annotation of probes was performed with R/Bioconductor package illumina HumanMethylation450kdb, with mapping to the hg19 genome build. Enriched gene ontology (GO) pathways were by interrogation of the annotated genes with significant CpGs with DAVID (Huang da *et al.*, 2009). Differentially methylated regions (DMRs) were identified by R/Bioconductor package DMRcate. DMRs are defined as regions containing >1 differentially methylated CpG with a maximum separation of 1000bp.

### Integration with gene expression

Gene expression was profiled by illumina whole-genome expression array for Day0 and Day14 MSC chondrogenesis triplicate biological samples and expression analysis was performed in the R/Bioconductor Limma package (Barter *et al.*, 2015). For genes with multiple probes gene expression values were averaged then intersected with DNA methylation Beta values associated with gene loci. Correlation of significant CpGs with gene expression was performed on binned log2 chondrogenesis gene expression changes to identify the percentage CpGs in at gene loci with methylation changes.

### Chromatin states

To assign MSC-derived chondrocyte chromatin states to probes, we downloaded the NIH Roadmap Epigenomics Mapping Consortium E049 15-state chromHMM model and mapped probes based on their locations in the genome (Roadmap Epigenomics *et al.*, 2015) (http://www.roadmapepigenomics.org/). Roadmap chromatin states were collapsed into five functional states: Transcription Start Site (TssA, TssBiv), Flanking Transcription Start Site (TssAFlnk, BivFlnk), Transcribed (Tx, TxFlnk, TxWk), Enhancer (Enh, EnhG, EnhBiv), and Repressed (ZNF/Rpts, Het, ReprPC, ReprPCWk, Quies). All plots were generated using the ggplot2 package in R.

### Comparison with articular chondrocyte and TCGA normal tissue 450K methylation data

Publicly available 450K DNA methylation profiles were downloaded for human articular cartilage neck of femur fracture control (NOF) and osteoarthritic diseased (OA) (GEO GSE63695) and from The Cancer Genome Atlas (TCGA) for non-cartilage tissue healthy control samples (https://cancergenome.nih.gov/) (Rushton *et al.*, 2014). All tissue (cartilage, TCGA and Day0/14 MSC chondrogenesis) Beta values underwent quantile normalisation. Chondrocyte-specific hypomethylated CpGs were defined as CpG sites with an average methylation beta value <0.5 in NOF cartilage and >0.8 in >90% of TCGA non-cartilage tissues.

### Data availability

MSC chondrogenesis 450K methylation data is deposited at…

---

Supplementary Figure 1.

Supplementary Figure 2.

Supplementary Figure 3.

Supplementary Table 1.

DML

Supplementary Table 2.

DMR

Supplementary Table 3.

MSC and OA CpG

